# Proteomic profile analysis of plasma and aqueous humor from glaucoma and non-glaucomatous patients

**DOI:** 10.1101/2024.04.11.588885

**Authors:** Carmen L. Pessuti, Chia-Ling Huang, Angela Banks, Nhi Vo, Lori Jennings, Joseph Loureiro, Kleber S. Ribeiro, Deise Fialho Costa, Heloisa Nascimento, Cristina Muccioli, Ivan Maynart Tavares, Alessandra G. Commodaro, Rubens Belfort, Christopher W. Wilson, Amy Chen, Ganesh Prasanna, VijayKrishna Raghunathan

**Author notes:** **Corresponding author(s):** VijayKrishna Raghunathan, Ph.D., Ophthalmology, BioMedical Research, Novartis, 22 Windsor St, Cambridge, MA, Massachusetts, 02139. *Email:*; Ganesh Prasanna, Ph.D., Ophthalmology, BioMedical Research, Novartis, 22 Windsor St, Cambridge, MA, Massachusetts, 02139. *Email:.

## Abstract

**Purpose:** Glaucoma, a multifactorial ocular neuropathy, can lead to irreversible vision loss. Diagnosis involves assessing optic cupping (increased cup-to-disc ratios) and structural changes (like retinal nerve fiber layer thinning) through clinical imaging. Elevated intraocular pressure (IOP) is commonly associated with glaucoma, but not always. However, understanding disease progression is hindered by limited access to donor ocular tissue and consistent clinical data. We hypothesize that the proteome of aqueous humor and plasma may be altered in disease and correlates with clinical parameters such as IOP and cup-to-disc ratios.

**Methods:** Aqueous humor (AH) and plasma samples were collected from 36 glaucoma patients (17 male, 19 female), and 35 non-glaucomatous control patients (16 male, 19 female) undergoing cataract surgery. The protein profile was compared using the SOMAscan® assay system for proteome profiling. From glaucomatous donors, correlations between IOP and cup-to-disc ratios to proteome differences were determined.

**Results:** Overall proteomics profiles between both AH and plasma were compared by combining all samples (glaucoma and non-glaucoma) and then performing correlation analyses. This study revealed similar protein abundance in the two biological fluids. Additionally, it identified different abundance of proteins in plasma and AH between glaucoma and non-glaucoma samples. The differential proteins identified were involved in pathways related to vascular integrity, inflammation, immune response, cell adhesion, and complement activation. Generally, glaucomatous AH showed higher protein levels. Neurofilament light chain (NEFL) protein correlated with elevated IOP and inflammatory markers, but not with cup-to-disc ratios.

**Conclusions:** Together, our data demonstrate that the proteins identified in this study from glaucomatous donors correspond to markers of neurodegeneration and those that may inhibit cell proliferation or disrupt vascular integrity.

## INTRODUCTION

Primary open angle glaucoma (POAG) is one of the leading causes of irreversible blindness, estimated to affect 111.8 million people worldwide by 2040 ^1,2^. Although the etiology and progression of POAG is multifactorial, the only modifiable causal risk factor for managing visual field loss is to reduce intraocular pressure (IOP), which is demonstrated to slow the progression of damage to the optic nerve and death of retinal ganglion cells ^3^. IOP is largely determined by a balance between aqueous humor production and resistance to outflow ^4,5^. In the human eye, approximately 75% of all aqueous humor flows from the anterior chamber through the trabecular meshwork (TM) and Schlemm’s canal (SC) ^6,7^ and is thus the primary egress for aqueous humor ^8^. Increased resistance to aqueous drainage due to changes in TM cells and extracellular matrix (ECM) is considered a major contributor to ocular hypertension (OHT) ^9,10^. Differential profiles and levels of soluble factors have been identified in the AH, which have the potential to affect cells in the tissues (outflow tract) that they traverse through, and thus influence OHT.

AH is secreted into the posterior chamber by the non-pigmented epithelial cells of the ciliary processes. Though the total amount of protein in the anterior chamber is less than 1% (w/v) due to the functional blood aqueous barrier ^4^, the aqueous humor is enriched with proteins secreted by intraocular structures from both the anterior and posterior segments ^11^. It is thus reasonable to hypothesize that systemic disorders, genetic conditions, or perturbations to ocular integrity and health may all adversely influence expression of proteins in the aqueous humor and other ocular tissues ^12^. Disease associations of protein compositions of tear films, AH and vitreous humors have been observed with dry eye, glaucoma, age related macular degeneration, or other pathologies ^13–18^, although very few studies have correlated these with clinically relevant parameters.

Several studies have demonstrated differences between metabolites or proteomes in the AH of patients with primary open angle glaucoma and have subsequently posited that these may serve as biomarkers of dysfunction in the outflow apparatus ^16,19–29^. Proteomic analysis of aqueous humor reveals significant protein expression differences between cataract and primary open angle glaucoma (POAG) patients. Multiple studies report that dozens to hundreds of proteins are differentially expressed in POAG, involving pathways such as oxidative stress, inflammation, lipid metabolism, extracellular matrix regulation, and neural degeneration ^30–41^. Key findings from these various studies identify several proteins in the complement cascade, apolipoproteins, and antioxidant defense suggesting immune involvement and metabolic stress in POAG.

For the aforementioned investigations, complementary approaches were used to quantify protein expression in AH from glaucomatous donors, including ELISA, multiplexed immunoassays, LC-MS/MS, or antibody microarrays. Limitations in quantifying proteins/metabolites include the small volume and low protein concentration in AH, alongside the need for validated antibody-based methods, which restricts biomarker identification. Thus, more sensitive methods are needed to broadly identify AH proteins to find disease progression biomarkers when comparing pre- and post-treatment stages. Discovering new proteins in the AH of patients could indeed shed light on the molecular mechanisms behind outflow resistance in OHT and glaucoma. Overall, while these proteomic changes provide insight into POAG mechanisms and support biomarker development, limited studies^42^ have performed direct comparisons between plasma and aqueous humor or with clinical parameters such as IOP or cup-to-disc ratios. Others have suggested that differences in proteome profiling within POAG populations may enable subgrouping to identify POAG severity^43^, though these studies lacked clinical correlation. We hypothesize that proteins secreted into the AH and plasma of patients with glaucoma differ from non-glaucomatous individuals and are associated with clinically relevant parameters. Here, we utilize the SOMAscan® assay ^44^, a highly multiplexed, aptamer-based technology, to detect and assess relative abundance of a broad range of proteins from small volumes of human blood alongside AH from patients undergoing glaucoma surgery, as compared to controls patients subject to cataract surgery.

## MATERIALS AND METHODS

### Study population

The study was approved by the ethics committee investigational review board (CEP 1.209.725) and adhered to the principles of the Declaration of Helsinki. Informed consent was obtained from all participants. Aqueous humor (AH) and plasma samples were collected from 32 glaucoma patients (16 male, 16 female), and 35 cataract control patients (16 male, 19 female) undergoing cataract surgery (**Figure 1**). The control cohort underwent clinical examinations (e.g. elevated IOP, optic nerve head changes, or uveitis) to rule out any glaucoma-related findings as part of inclusion/exclusion criteria. Within the glaucoma patient group one underwent trabeculectomy, two were pseudophakic, and AH was collected at the beginning of the surgery. Clinical diagnosis of glaucoma was based on elevated IOP (IOP > 21 mmHg), documented progressive cupping or thinning of the neuroretinal rim. All glaucoma patients were treated with medication to control IOP. Patients with other eye diseases or associated with systemic diseases were not included. Our study is exploratory in nature, and the primary focus is on the proteomic analysis of plasma and aqueous humor samples from the study participants. As such, the data about the glaucoma patients in this study will be limited to the existing inclusion criteria and general demographic information. Additional longitudinal clinical data including summary statistics of HVF, glaucoma phenotype, number of medications, etc. were unavailable and thus not being reported here for this study. All donor meta data including IOP, cup to disc ratios and visual acuity are included as **Supplementary Table 1**.

**Figure 1:**
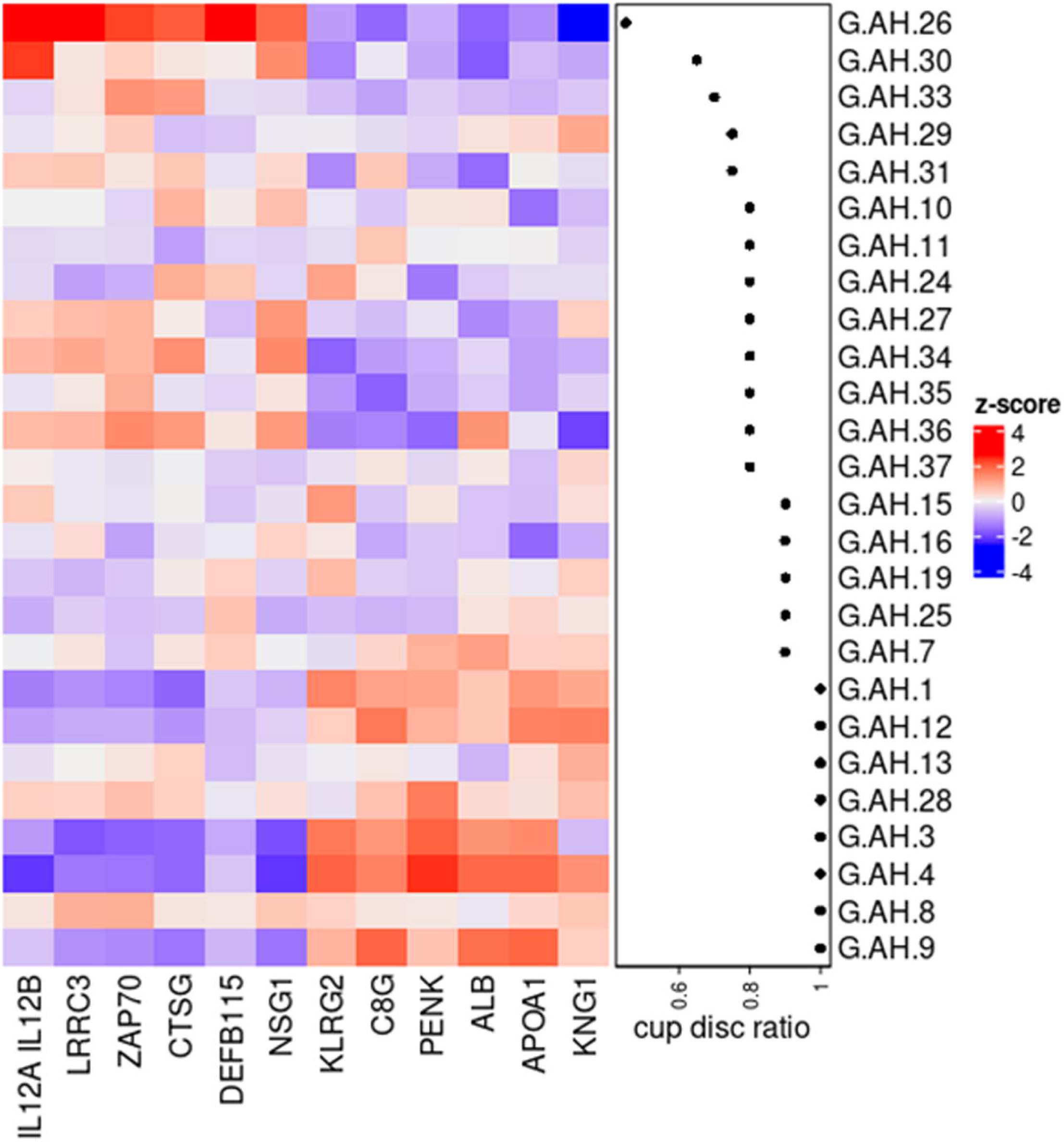
Patient demographics demonstrate study participants were age and sex matched. Intraocular pressures and protein concentrations from AH are reported for both glaucomatous and non-glaucomatous donors.

### Aqueous humor (AH) and plasma sample collection

AH samples were collected from patients following a previously validated method ^45^ at the beginning of the surgery. Briefly, after anesthetic and antiseptic eyedrops administration, a 1mL tuberculin syringe (27 gauge, ½ inch) was used to aspirate the AH. Approximately, 0.1 ml was collected, and transferred to two labeled Eppendorf tubes. Peripheral blood (10 mL) was collected from an arm of the patient in EDTA tubes during the surgery. The tubes were centrifuged for 10 min at 1900 x g, and plasma was collected. Total protein concentrations in the AH samples were determined using the BCA Protein Assay Kit (Pierce, Rockford, IL).

### SOMAscan® analysis

All AH and plasma samples were analyzed in parallel using the SOMAscan® proteome profiling platform that provides a broad and relative abundant assessment of selected proteins in biological fluid. The SOMAscan® platform, using affinity-based slow off-rate modified aptamers (SOMAmers), has been described extensively elsewhere ^44,46,47^. The custom SOMAscan® version used in our study covered 5,080 SOMAmers, including 4,785 human-specific SOMAmers targeting 4,161 human proteins. Evidence supporting SOMAmer endogenous antigen annotation, including SOMAmer validation by proteogenetic evidence, mass spectrometry detection, or orthogonal assay concordance, where known, are provided in the referenced studies^48–66^. The targeted proteins include transmembrane receptors, secreted proteins, kinases, transcription regulators and signal transducers. The samples (55 µl) were aliquoted into 2D barcoded tubes and were shipped on dry ice to SomaLogic for analysis.

### Statistical analysis

The relative abundance of captured SOMAmers were quantified by microarray hybridization as relative fluorescence units (RFU). The annotations for the SOMAmers and log_2_(RFU) for all samples analyzed are provided in **Supplementary table 2**. Quality control and sample outlier detection was conducted using the arrayQualityMetrics^67^, and by running principal component analysis (PCA) and Spearman’s correlation analysis with custom R scripts. RFU were log2- transformed and normalized by smooth quantile normalization using R package qsmooth ^68^. In total, 5 samples (2 non-glaucomatous, 3 glaucoma) were removed from the AH cohort and 2 non-glaucomatous samples were removed from the plasma cohort, which left 62 samples in the AH cohort and 65 samples in the AH cohort. Exclusion of 5 samples were based on an unbiased statistical outlier analysis (**Supplemental Figure 1**) and are thus not presented in subsequent analysis. We applied the limma R package (v3.50.3) in R 4.1.0 environment to determine differentially expressed proteins between groups^69^. Pathway enrichment analysis was performed with pre-ranked GSEA using R package fgsea v1.20.0 and Gene Ontology Biological Process category (GOBP) in MSigDB^70–73^.

### Correlation analysis

We conducted Elastic Net, a machine learning based method, to identify proteins correlated with the total protein concentration. For this analysis, we utilized the cv.glmnet function from the R package glmnet v4.1-3. We employed a 10-fold cross-validation approach to optimize the alpha and lambda parameters^74,75^. Additionally, we performed a 1000-fold bootstrapping using the optimized alpha and lambda values to determine the frequency at which each protein was selected as a top predictor for estimating the total protein concentration. To enhance the performance of our analysis, we only included proteins with an absolute Pearson’s correlation coefficient greater than 0.4 with the total protein concentration.

## RESULTS

### Donor demographics

Donor demographics for the patient samples used in this study are provided in **Figure 1**. The average age of donors with cataract was 66.9 ± 10.9 while those with glaucoma was 68.41 ± 12.84. No significant differences (p=0.60, t-test; **Figure 1**) were seen in donor ages between the groups *in toto* or when evaluated by sex. Amongst these donors, we note that one donor had primary angle closure glaucoma while the remainder had POAG. The mean IOP of glaucoma donors (IOP controlled with medication) was 17.94 ± 8.28 mmHg with no differences seen between male or female donors. Only one POAG donor had an IOP > 24 mmHg. A statistically significant difference in total protein concentration in AH was observed between cataract and glaucoma groups (Cataract: 0.383 ± 0.505 mg/ml; Glaucoma: 1.17 ± 1.48 mg/ml). Visual acuity and cup-to-disc ratios of POAG donors are listed in donor metadata within **Supplementary Table 1**.

### Differential analysis of SOMAScan® profiling data

Principal component analysis of the cataract and glaucoma samples revealed heterogeneity between samples, including outliers that were subsequently excluded from further analysis. Subsequent analysis revealed few differences in protein levels between non-glaucomatous and glaucomatous aqueous humor and plasma (**Supplementary Fig. 1**). Nevertheless, we observed that samples were clustered by tissue types with reasonable correlation (r > 0.5) between AH and plasma samples (**Figure 2a**). The histogram (**Figure 2b**) demonstrates that the distribution of Pearson’s correlation coefficients between AH and plasma samples from same donor demonstrates some variability. Considering all donors, a reasonable correlation (r > 0.5) was observed comparing protein levels between AH and plasma samples.

**Figure 2:**
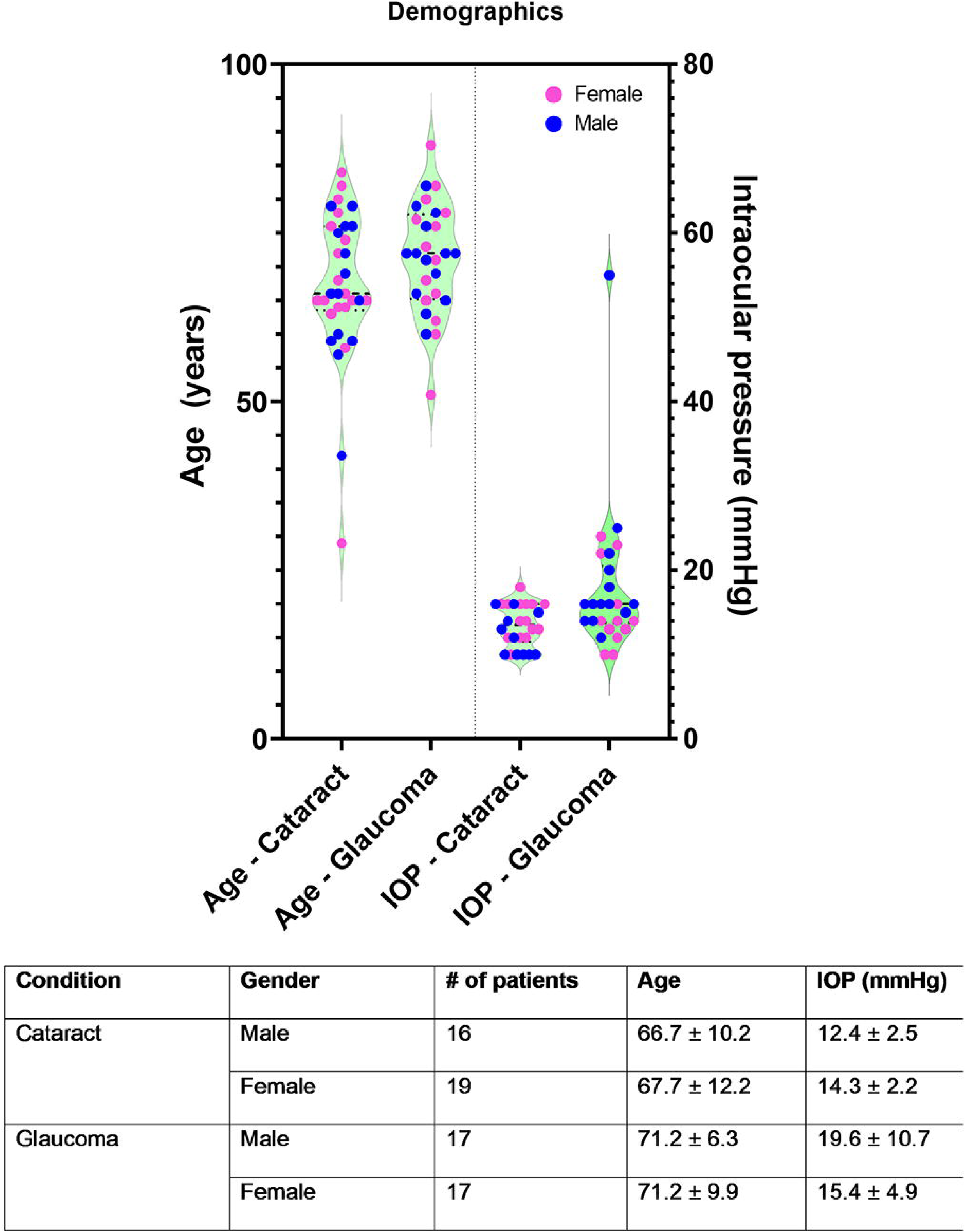
Proteins identified in aqueous humor and plasma clustered by type of biofluid, and appear to correlate reasonably well as evidenced by Spearman’s correlation. **(A)** Heatmap, and **(B)** histogram of Spearman’s correlation index demonstrates that the correlation, while evident, is not absolute i.e. 0.5 < R^2^ < 0.7.

Subsequent to multivariate analysis, we performed differential protein analysis for both plasma and aqueous samples. Expression values that were |log2(RFU)| > 0.5 and where the adjusted p-value was < 0.05 are highlighted as significant (**Figure 3**). Within glaucomatous AH (Figure 3A), 7 SOMAmer probes representing 6 target proteins were observed to be significantly down regulated, while 49 SOMAmer probes representing 49 target proteins were upregulated. Examples of top up-regulated proteins include complement proteins (C3, C7, C1QTNF3), Apolipoprotein (APOE, APOF), matrix proteins (endostatin/Col18A1), macrophage associated proteins (CSF1R, CD163), while top down-regulated proteins include growth factors such as FGF9, FGF20 and CCK. We observed that in plasma samples, the majority of the differential proteins were upregulated in glaucoma. Some of the upregulated proteins of interest include ENO1, NEK7, AKT, PRKCA, PRKCB, JAK2, MAP kinase-activated protein kinase 2 (MAPKAPK2), whereas the top down-regulated proteins include APOC3, GDF2, DCP1B and EDIL3. A comprehensive list of proteins altered in their levels comparing glaucomatous and non-glaucomatous plasma and AH are provided in **Supplementary table 3**. Although protein levels from AH and plasma from the same donor showed a positive correlation, we only observed SNX4 to be significantly up regulated in both AH and plasma samples from glaucomatous donors.

**Figure 3:**
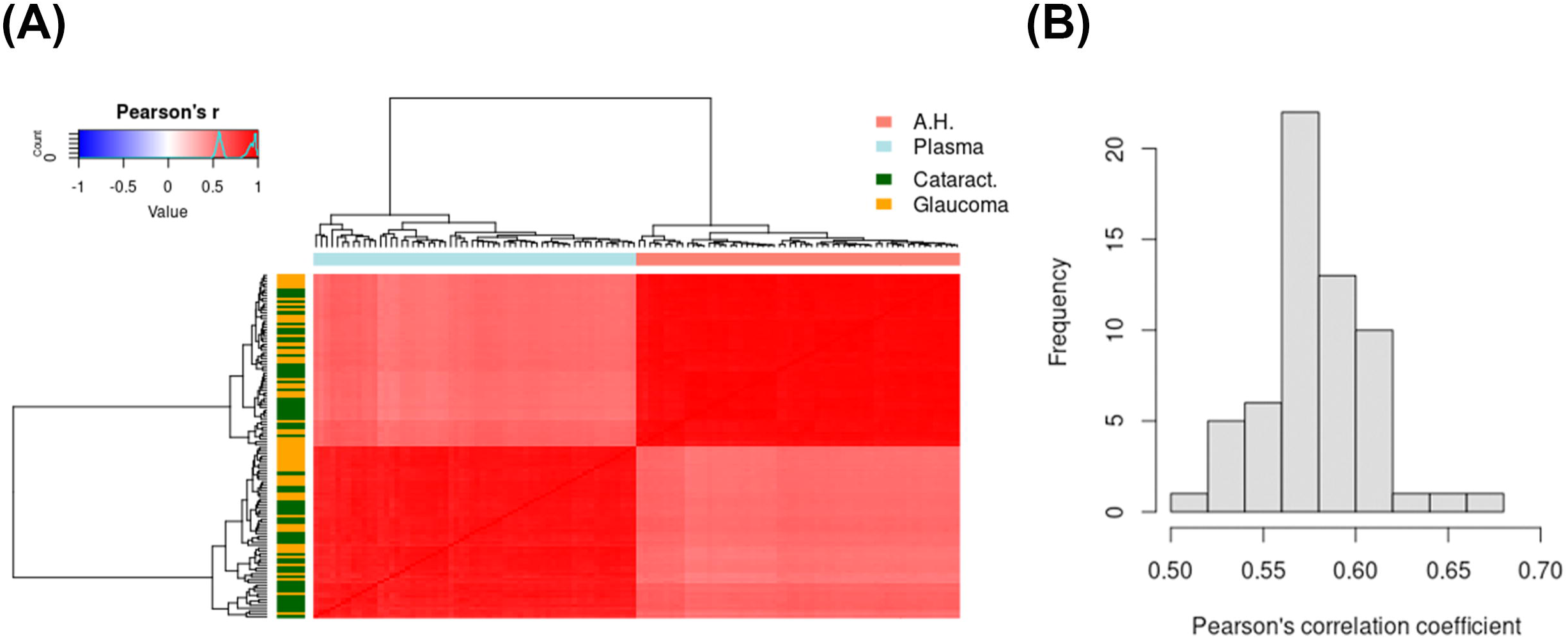
Principal component (PCA) and differential expression analysis demonstrates differences in proteins identified in **(A)** aqueous humor and **(B)** plasma comparing glaucomatous and non-glaucomatous donors. PCA demonstrates large variance between cataract (non-glaucoma) and glaucomatous donor samples with no discernible separation between the two disease groups. Volcano plots demonstrate that no significant differences were seen comparing cataract (non-glaucoma) and glaucomatous donors for most proteins. Few proteins whose levels were significantly altered are represented in the bar plot.

### Pathway enrichment analysis

Next, we asked if the differentially enriched proteins were enriched in common biological pathways, by performing pathway analysis (**Supplementary table 4**). We identified 51 and 559 significantly up-regulated pathways (adjusted p-value < 0.05) in AH and plasma samples from glaucoma donors, respectively. 16 pathways were up-regulated in both AH and plasma, including inflammatory response, IL6 production and immune response related pathways.

### Correlation analysis of protein levels with total protein concentration

While individual protein changes may help identify signaling pathways perturbed in disease, total protein concentration of the AH may help ascertain the overall health of the blood-aqueous barrier and the outcomes of IOP lowering medications. Thus, normalizing individual proteins to total protein concentrations assists in determining if observed changes reflect pathway changes underpinning disease, or are merely a consequence of structural failure or drug treatment(s). We conducted an Elastic Net regression analysis to identify proteins that can accurately predict the total protein concentration (**Figure 4a**). Our analysis specifically focused on AH proteins, as proteins from these samples may provide a better reflection of changes in glaucoma disease compared to plasma samples. Through our analysis, we successfully identified 5 proteins (KNG1, IGFBO6, MAP2K4, H6PD, and C3) that demonstrated the highest predictive capability for the total protein concentration collected from the patients (**Figure 4a**).

**Figure 4:**
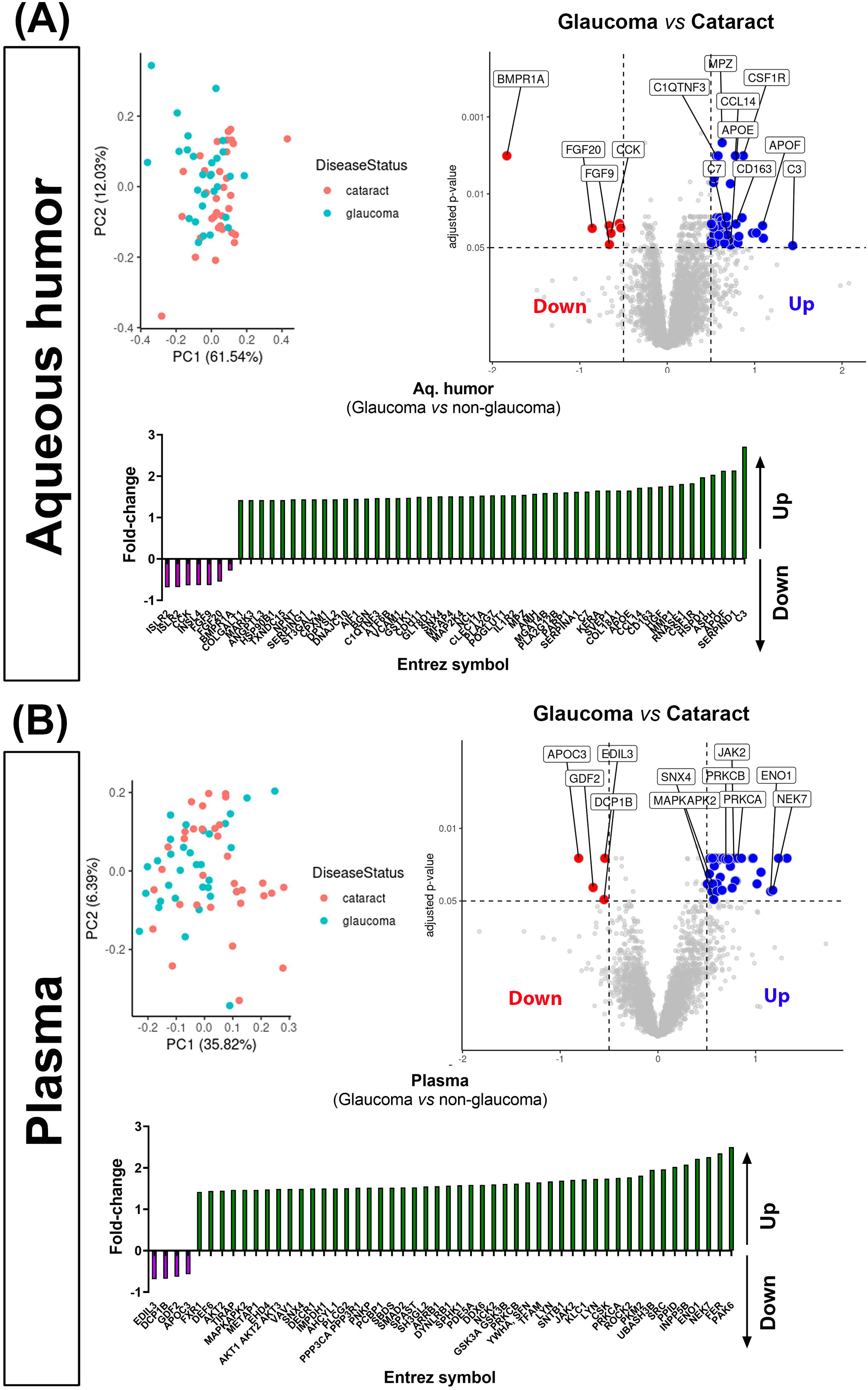
Elastic net regression analysis (aqueous humor only) demonstrated **(A)** increased protein concentration is observed in aqueous humor of donors with glaucoma, with no specific sex related differences observed, **(B)** changes in some proteins correlate significantly with IOP independent of sex of donor. **(C)** Scatter plots and linear regression of proteins (R^2^ > 0.25) as a function of IOP indicate NEFL, DLL4, and NFE2L1 to most significantly altered proteins in the aqueous humor.

### Correlation analysis of protein levels with IOP

We conducted Elastic net regression analysis to identify 13 proteins to be associated with IOP values reported in glaucomatous patients (**Figure 4b**). We note that a glaucoma sample with extremely high IOP measurement was excluded from the analysis. Among the proteins of interest in glaucoma, the protein with maximal correlation with IOP levels in glaucomatous patients was neurofilament light chain (NEFL; **Figure 4c, Supplementary** Fig. 2). Linear regression analysis validated this finding and demonstrated a net correlation with IOP measurements (R^2^ = 0.3067). Since NEFL is a widely accepted marker for several neurodegenerative diseases including glaucoma ^76–85^, we then decided to further investigate if levels of NEFL correlated with other SOMAmers identified in glaucomatous donors. We calculated the Pearson’s correlation coefficient of NEFL and rest of the proteins and identified 29 proteins that are positively or negatively correlated with NEFL (|r > 0.4|), including TEAD3, CNTF, C4A, C4B and CCL27 (**Figure 5**).

**Figure 5:**
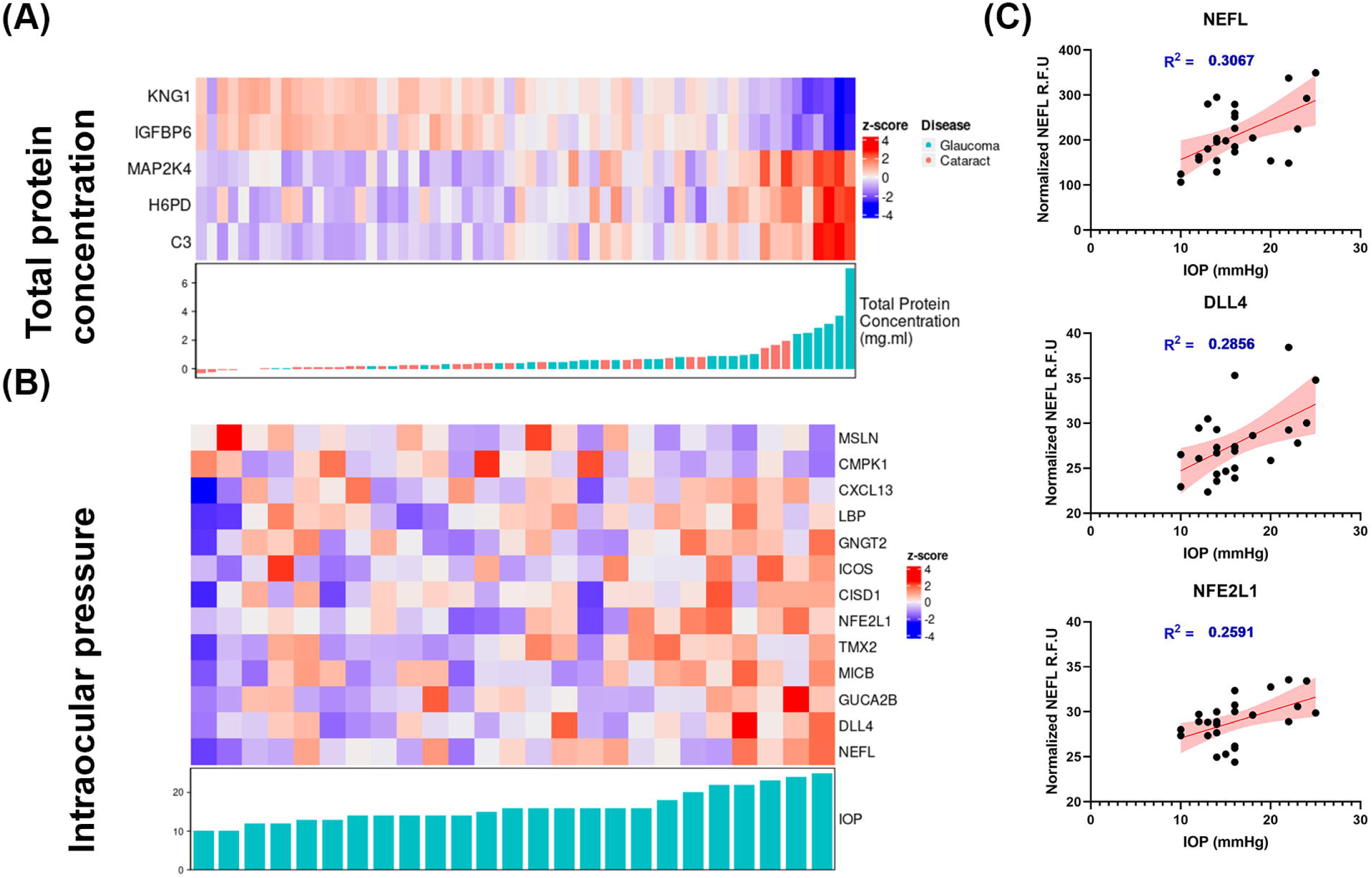
Heatmap demonstrates correlation of NEFL to top 30 proteins identified in the aqueous humor of glaucomatous donors. Pearson’s correlation analysis comparing NEFL with rest of the proteins identified ∼30 proteins that either positively (red) or negatively (blue) correlated with NEFL (|r > 0.4|). In general, more proteins that negatively correlated with NEFL levels were identified than those that exhibited positive correlation.

### Changes in protein levels as a function of cup-to-disc ratios in glaucoma

Finally, we sought to determine if changes in relative abundance of protein levels in AH correlated with cup-to-disc ratios of glaucomatous patients via Pearsons’s correlation analysis (**Figure 6**). For patients where cup-to-disc ratios were available for both eyes, we used the average measurements from both eyes. We note that among the 29 glaucomatous samples obtained, cup-to-disc ratios were available for 26 donors. Figure 6 shows the top 6 positive (KLRG2, C8G, PENK, ALB, APOA1, and KNG1) and negative (IL12A/B, LRRC3, ZAP70, CTSG, DEFB115, and NSG1) correlated proteins with cup-to-disc ratios.

**Figure 6:**
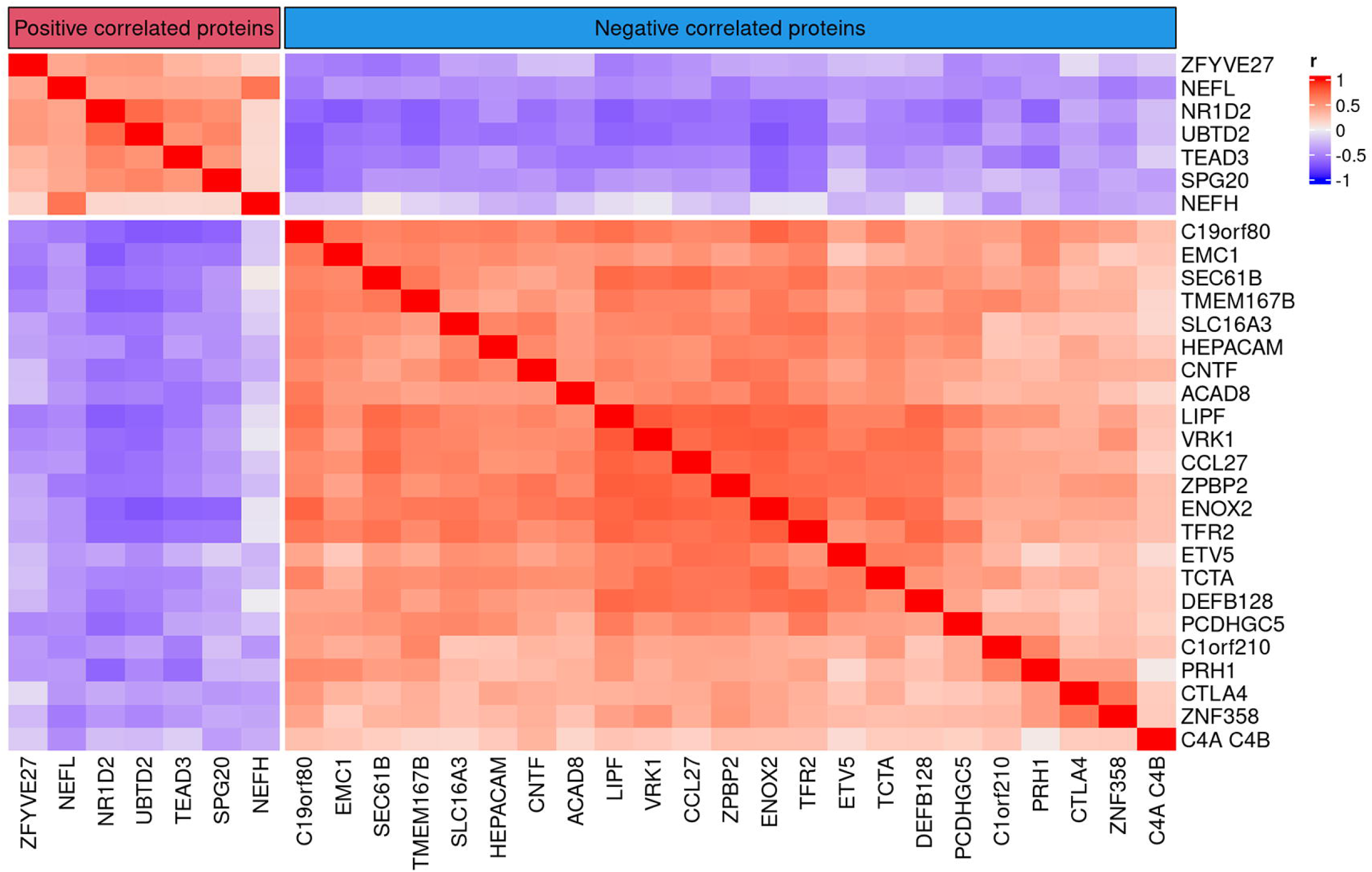
Heatmap identifies top 6 proteins either up or down regulated in the aqueous humor of glaucomatous donors as a function of cup-to-disc ratios.

## DISCUSSION

Monitoring glaucoma in the clinic typically involves measuring IOP, gonioscopy, and advanced imaging of the retina/optic nerve head, which require regular visits and patient compliance. Without a detailed medical history, it is challenging to understand the molecular mechanisms of disease progression or identify biomarkers to ascertain treatment effectiveness. Since collecting intraocular tissues from living patients is not feasible, analyzing biofluids such as AH and blood can provide critical insights into disease mechanisms. In this study, we examined protein levels in AH and plasma from patients with and without glaucoma undergoing cataract surgery using a SomaLogic platform and correlating these with IOP and cup-to-disc ratios measured immediately prior to the surgery.

### Comparative analysis of plasma and aqueous humor proteomes

The ages and sex of glaucomatous and non-glaucomatous donors in this dataset were comparable. We observed significant changes in relative abundance of a subset of proteins identified both in AH and plasma. Protein levels from AH and plasma from the same donor sample showed a positive correlation (r > 0.5 for all of samples). Interestingly, when comparing protein levels and correlations across all fluids and disease states, we only observed a significant upregulation of SNX4 in glaucomatous AH and plasma when compared with those from non-glaucomatous donors. SNX4 is a synaptic & endosomal membrane trafficking protein in the secretory pathway and is also implicated as a mitochondrial recycling quality protein in aging ^86–88^. The lack of greater correlation between plasma and AH proteins may be due to several reasons: turnover rates of AH *vs* plasma, tissues & cell types involved in protein secretions, systemic *vs* local transport of proteins. It is yet unclear if glaucoma is purely a disease of the eye or if systemic factors exist that regulate its etiology and progression, although co-morbidities have been reported ^89–93^. Further, Miguel Coca-Prados et al^94^ have identified several unique proteins and peptides in the AH attributed to the neuroendocrine nature of ciliary non-pigmented epithelial cells in addition to plasma proteins. It is thus unsurprising that differences in uniquely identified proteins exists between the two biofluids. Nevertheless, in the plasma, 48 proteins were upregulated while 4 (EDIL3, DCP1B, GDF2, APOC3) were downregulated when comparing non-glaucomatous and glaucomatous donors. Gene ontology analyses of proteins altered in the plasma of glaucomatous donors were related to multiple pathways corresponding to protein & macromolecule localization, intracellular transport, kinase activity pathways, post translational / post transcriptional regulation of proteins / genes, cytoskeletal reorganization and GPCR related pathways. On the other hand, 7 (ISLR2, CCK,

INSL4, FGF9, FGF20, BMPR1A) proteins were significantly downregulated and 49 were significantly upregulated in the AH of glaucomatous donors. Gene ontology analyses of proteins identified as altered in the AH of glaucomatous donors were related to multiple pathways corresponding to inflammatory response, humoral & adaptive immune response, cell-cell & cell-matrix adhesion, complement activation, and wound healing responses. These together suggest that though overall concordance in proteins may be observed between AH and plasma, some differences exist and thus proximity of biofluid to relevant tissue being studied (i.e. ciliary process, ciliary muscle and trabecular meshwork/Schlemm’s canal) may be important to consider. As such, deeper analysis in this study was focused on the ocular specific biofluid (AH) proteins.

### Effect of disease state on aqueous humor proteome

All changes in protein levels were normalized to total protein concentration and as such these changes are independent of mean protein concentration. Interestingly, regression modeling demonstrated that total protein concentrations trended towards being significantly greater in AH of glaucomatous compared with non-glaucomatous donors. While this is suggestive of a breakdown in the blood-aqueous barrier ^4^, whether this is due to the chronic nature of the disease or due to IOP lowering medications (e.g. prostaglandin analogs), other systemic medications or other co-morbidities is unclear. Nevertheless, the changes in proteins appear to be directly related to pathways with the potential for consequential effects on IOP regulation and outflow homeostasis. When comparing our overall findings with that from previous publications using other technologies, it was noteworthy that proteins related to complement and immune-related proteins (e.g. C3b, IL1, C1Q, C7 etc), apolipoproteins (APOE, APOF etc), heat shock/oxidative stress (HSPD1/Hsp60), matrix and serpin (SERPINA1, SERPING1, BGN etc) remained consistently differentially expressed between glaucomatous and non-glaucomatous AH^34,41,95–101^. These suggest that the proteomics methodology used in our study is sensitive and robustly identifies certain proteins, in glaucoma, regardless of study design or detection method. This highlights that certain pathways and proteins may have a functional or disease-correlative role that required further studies to establish their role in disease mechanisms. Nonetheless, several unique proteins were also identified in our study.

For example, Col18A1, which was upregulated in AH of glaucomatous donors, is expressed ubiquitously in ocular tissues except in photoreceptors, and protein fragments that contain endostatin (from Col18A1 cleavage) has been observed to accumulate in ocular fluid samples^102^. Interestingly, endostatin has been reported to promote expression and release of thrombospondin-1, implicated in outflow resistance & glaucoma ^103–105^, in non-ocular endothelial cells ^106^. Conversely, endostatin has also been posited to crosstalk with Rho/ROCK, TGFβ, NF- κB, PDGF, and autophagy pathways to yield anti-fibrotic effects ^107^. SVEP1, another overexpressed secreted ECM protein in AH & a disease modifier allele for congenital glaucoma ^108^, is known to be a binding partner for TIE1 and can thus regulate signaling outcomes in lymphatics and vasculature ^109^. Since AH predominantly drains into the Schlemm’s canal, a unique vessel with both vasculature and lymphatic properties ^110^, studying SVEP1 in the context of glaucoma warrants further investigation. Of particular interest, VCAM1, CDH11, and BGN were other proteins that were overexpressed in the AH of glaucomatous patients. SVEP1, CDH11 and VCAM1 were all identified as genes relevant to POAG pathogenesis in a genome-wide meta-analysis across ancestries ^111^. CDH11 can interact reciprocally with fibronectin binding protein syndecan-4 to facilitate cell migration and adhesion, partake in EMT, and modulate proliferation ^112–115^. In non-ocular cells, CDH11 was shown to mediate adhesion of macrophages to fibroblasts promoting transdifferentiation into myofibroblasts and a self-sustaining profibrotic niche ^116^. Biglycan (BGN) is an extracellular proteoglycan, expressed in the trabecular meshwork ^117^, is posited as a prognostic marker for cancer aggressiveness ^118^, fibrosis & inflammation ^119^ and is predicted to be a circulating “messenger” for triggering inflammation and/or autophagy ^120^. Whether biglycan in the AH serves as a biomarker, a signaling molecule to trigger autophagy or inflammatory phenotypes in the trabecular meshwork to mediate subsequent outflow regulation requires mechanistic studies. High levels of circulatory VCAM1 in the blood/plasma was found to correlate with ventricular hypertrophy, hypertension and is suggested to thus serve as a predictive soluble biomarker for cardiovascular disease and inflammation ^121–123^. In this study, whether VCAM1 expression is a result of chronic elevated IOP, drugs to lower IOP, or due to undiagnosed/unknown underlying systemic vascular conditions is unknown. Regardless, increased presence of the aforementioned proteins may serve as ‘proteins-of-interest’ in mechanistic understanding of outflow regulation for as potential biomarkers for dysregulation in outflow homeostasis. Further studies and orthogonal methods are needed to validate these findings.

### Protein changes identified as a function of clinical parameters in glaucoma

Since proteins with associations with ‘hypertension’ were identified, we next sought to determine if protein changes were a function of IOP levels through additional regression modeling. Two proteins notably were found to be of importance: DLL4 and NEFL. DLL4, a Notch ligand, is a key regulator of vascular morphogenesis, vessel maturation, and function. Secreted/soluble DLL4 has previously been reported to significantly reduce hydraulic conductivity, vascular permeability, and disrupt endothelial barrier function in non-ocular vasculature ^124,125^. Further, DLL4 has been demonstrated to be critical in developing retinal vasculature, increased endothelial cell proliferation, and angiogenic sprouting ^126^. Interestingly, DLL4 was also shown to inhibit inflammatory choroidal neovascularization despite opposing effects seen in endothelial cells (anti-angiogenic) and macrophages (pro-angiogenic) ^127^. Collectively, we speculate that DLL4, found upregulated with IOP, may act to increase outflow resistance via a yet unknown mechanism. Further, whether a dichotomous function for the DLL4 ligand or the Notch pathway exists in outflow regulation remains to be seen. A recent study suggests that Notch pathway protein expression may differ between segmental flow regions of healthy and glaucomatous donors, at least *in vitro* ^128^. Thus, ligands circulating in the aqueous humor could differentially impact TM cell function necessitating further studies.

NEFL is a well-established marker of neurodegeneration ^77,79–82,129^ and similar to our finding, has previously been reported to be elevated in glaucomatous AH in humans and animal models ^76,78,130^. In fact, NEFL had the highest correlation with IOP in all donors in this study, providing additional confidence that NEFL in the AH may indeed be a suitable biomarker for glaucomatous neurodegeneration and ocular hypertension. However, we advise caution that AH samples utilized in this study were from patients identified for glaucoma filtration surgery. Therefore, it is likely that the disease stage may be advanced and thus markers of axonal degeneration are expected within the AH. Whether NEFL would serve as an early marker of disease is unclear and thus requires comprehensive longitudinal natural history investigations. Further the source of the neurofilament protein, determined in the aqueous humor, is unclear (i.e. whether it is due to degeneration of the optic nerve, secretion from cells of neural crest origin (developmentally) in the anterior segment, or Schwann/nerve cells in the anterior segment remains unknown). Interestingly, while NEFL levels correlated with IOP, its levels did not appear to correlate with cup-to-disc ratios of glaucoma donors (**Supplementary Fig. 3**), though other proteins that did correlate were identified. Interestingly, cup-to-disc ratio was independent of IOP values as well (**Supplementary Fig. 3**). Inflammatory proteins (such as interleukins and cathepsins) corresponded with low cup-to-disc ratio, while apolipoprotein A1 (APOA1), albumin, kininogen-1 (KNG1) and complement C8G correlated with a higher cup-to-disc ratio suggesting proteins altered in the AH may differ based on disease severity. KNG1 is an antiangiogenic molecule that has been suggested to be a marker of neurodegeneration ^131^ and can impair the proliferation of endothelial cells ^132^. To the best of our knowledge, the role of KNG1 in POAG and/or ocular hypertension has not been studied, although its cleaved nonapeptide, Bradykinin (BK), has been the target for IOP lowering investigations ^133–138^. A prior study reported a decrease in C8G ^139^ in glaucomatous AH conflicting with our results. However, Kim et al ^140^ previously reported C8G to act as a neuroinflammation inhibitor; our result correlating C8G levels with high cup-to-disc ratios may thus reflect the advanced stage of neurodegeneration/neuroinflammation in glaucoma.

Consistent with our study, elevated levels of APOA1 were reported in AH of POAG donors^141^. It is important to note that the direct role of APOA1 in IOP homeostasis is not known.

However, APOA1 plays a critical role in reverse cholesterol transport pathway via direct interactions with the ABCA1 gene whose variants are implicated in POAG ^142^. Cholesterol is itself a risk factor for elevated IOP ^143^. Elevation in albumin levels in glaucomatous AH is also reported in several studies, though attributed to administration of IOP lowering drugs ^144^. Finally, a correlation analysis of NEFL to other proteins identified in glaucomatous aqueous humor revealed that NEFL levels may be associated with proteins regulating apoptosis, complement activation, proliferation, and cytoskeletal reorganization. Together, the proteins identified in this study correspond to both markers of neurodegeneration and those that may inhibit proliferation or vascular integrity.

## LIMITATIONS

This study is not without limitations. One of the major limitations is that the biofluids obtained in this study were obtained at the time of ocular surgery thus representing a singular snapshot in time and no information was gleaned about the stage of the disease; although, clinically measured IOP measurements and cup-to-disc ratios are available at this stage. We understand that glaucoma can be asymmetric, and in future studies, we will consider incorporating cup-to-disc ratios from objective approaches such as OCT to validate or refine these findings. Also, while cup-to-disc ratios are a clinical parameter, and protein levels (which potentially are dynamic) are a molecular parameter, it is important to note that these variables are independent, and that the kinetics of structural changes may differ from the kinetics of molecular turnover. The independent parameters were nevertheless compared to ascertain if the clinical parameter investigated had any association with the proteome content in the AH and if this could perhaps reflect the current state of the disease in the donor. Under the current study design, it is important to note that any causal or correlational relationship is hard to conclude since there are practical limitations on frequency of aqueous humor sampling. Further, comprehensive and longitudinal patient histories of systemic co-morbidities or ocular diseases and medications that may affect systemic or ocular hypertension including but limited to intraocular pressure were unavailable. Since IOP in glaucoma patients was controlled with IOP lowering medication, the impact of these therapeutics on proteome changes cannot be ascertained. Further functional measurements for visual field or additional structural deficits including longitudinal and pre-surgical fundus photography or OCT measurements (nerve fiber layer thinning, rim width, ganglion cell analysis etc.) are not known for these patients. We speculate that additional information that may be obtained from *longitudinal prospective studies* with disease state factored in could further enable identification of determining factors driving changes in the proteome. We did not perform any genetic linkage or association analysis to identify any polygenic risk assessment from the POAG patients enrolled. Future studies may be needed to identify correlations between genetic causes, structure-function changes, and molecular profiling approaches in a longitudinal manner if feasible. Since the entirety of the samples collected in this study were utilized for proteomics, attempts to validate the results using orthogonal or secondary methodologies were not undertaken. As such, since mass spectrometry (a commonly used profiling technique for proteomics) was not part of the current study’s scope, we recommend future research from independent investigators to include such methods, and to compare SOMAscan findings with previously published aqueous humor proteomic profiles and/or methods. Thus, we anticipate that the approach and data presented here will enable the design of validation studies for future investigations within the glaucoma community.

## ACKNOWLEDGEMENTS

The authors would like to thank the patient donors for the biological fluid samples without whose consent these experiments would be impossible.

## FUNDING STATEMENT

This work was sponsored by Novartis Biomedical Research, and CNPq (Ministry of Science, Technology, and Innovation of Brazil).

## AUTHORS’ DISCLOSURE

C.H, A.B, N.V, L.J, J.L, C.W.W, A.C, G.P, V.R are all employees of Novartis Biomedical Research.

## CONTRIBUTION STATEMENT

C.L.S, K.S.R, D.F.C, H.N, C.M, I.M.T, A.G.C, R.B.J: clinical design, sample collection, data interpretation, critical review, revisions, and approval of article. C.H, A.B: data analysis, data interpretation, critical review, revisions, and approval of article. A.C, G.P: Conceptualization of study, data interpretation, critical review, revisions, and approval of article. N.V, L.J, J.L: sample preparation & analysis, data analysis, data interpretation, critical review, revisions, and approval of article. C.W.W: data interpretation, critical review, revisions, and approval of article. V.R: data analysis, data interpretation, critical review, revisions, and approval of article.

## SUPPLEMENTARY INFORMATION

**Supplementary figure 1:** Outlier and principal component analysis identifying donors to exclude from data set in an unbiased manner.

**Supplementary figure 2:** Scatter plots of proteins identified in glaucomatous AH as a function of IOP demonstrate large variability between samples.

**Supplementary figure 3: (A)** Scatter plot of NEFL expression levels as a function of cup-to-disc ratios of glaucomatous donors demonstrate no significant relationship (R^2^ = 0.037) between the two parameters. **(B)** Cup-to-disc ratios of glaucomatous donors show no significant correlation (R^2^ = 0.008) with intraocular pressures reported.

**Supplementary table 1:** Donor metadata including age, sex, disease state, sample ID, IOP, cup-to-disc ratios & visual acuity.

**Supplementary table 2:** List of SOMAmer annotations and log_2_(RFU) for all samples analyzed from plasma and aqueous humor for both glaucoma and non-glaucoma donors.

**Supplementary table 3:** List of differentially expressed proteins in aqueous humor and plasma comparing glaucomatous and non-glaucomatous donors.

**Supplementary table 4:** GSEA pathway analysis of proteins identified in aqueous humor and plasma comparing glaucomatous and non-glaucomatous donors.

## Notes

### Competing Interest Statement

Several authors (VKR, GP, CW, AC, AB, CH, LJ, JL, NV) are employees of Novartis, BioMedical Research

### Summary of Updates

Raw data added as supplementary table Donor meta data added Limitations expanded Methods clarified

