## Supplementary figures and images for "Proteomic profile analysis of plasma and aqueous humor from glaucoma and non-glaucomatous patients"

### Fig S1

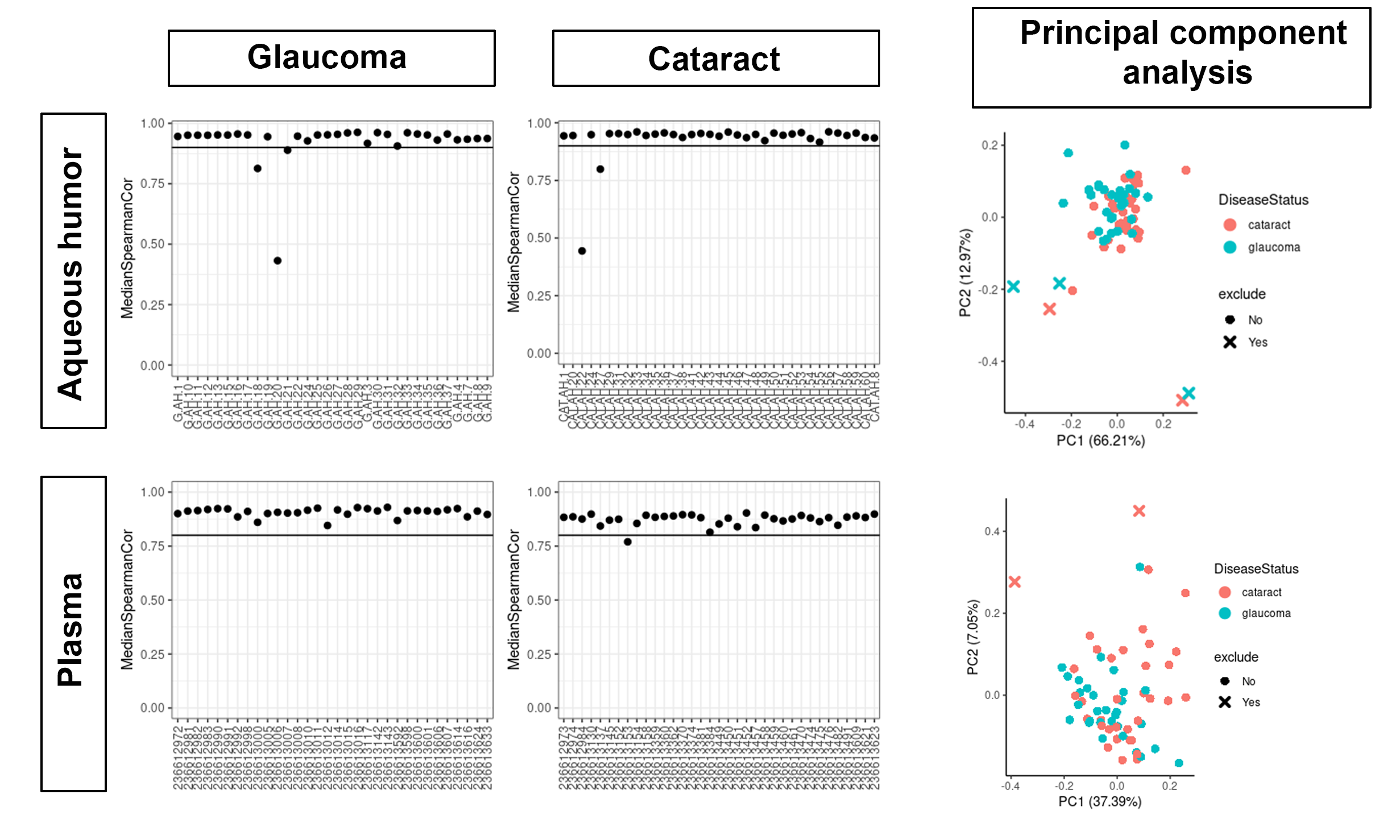

### Fig S2

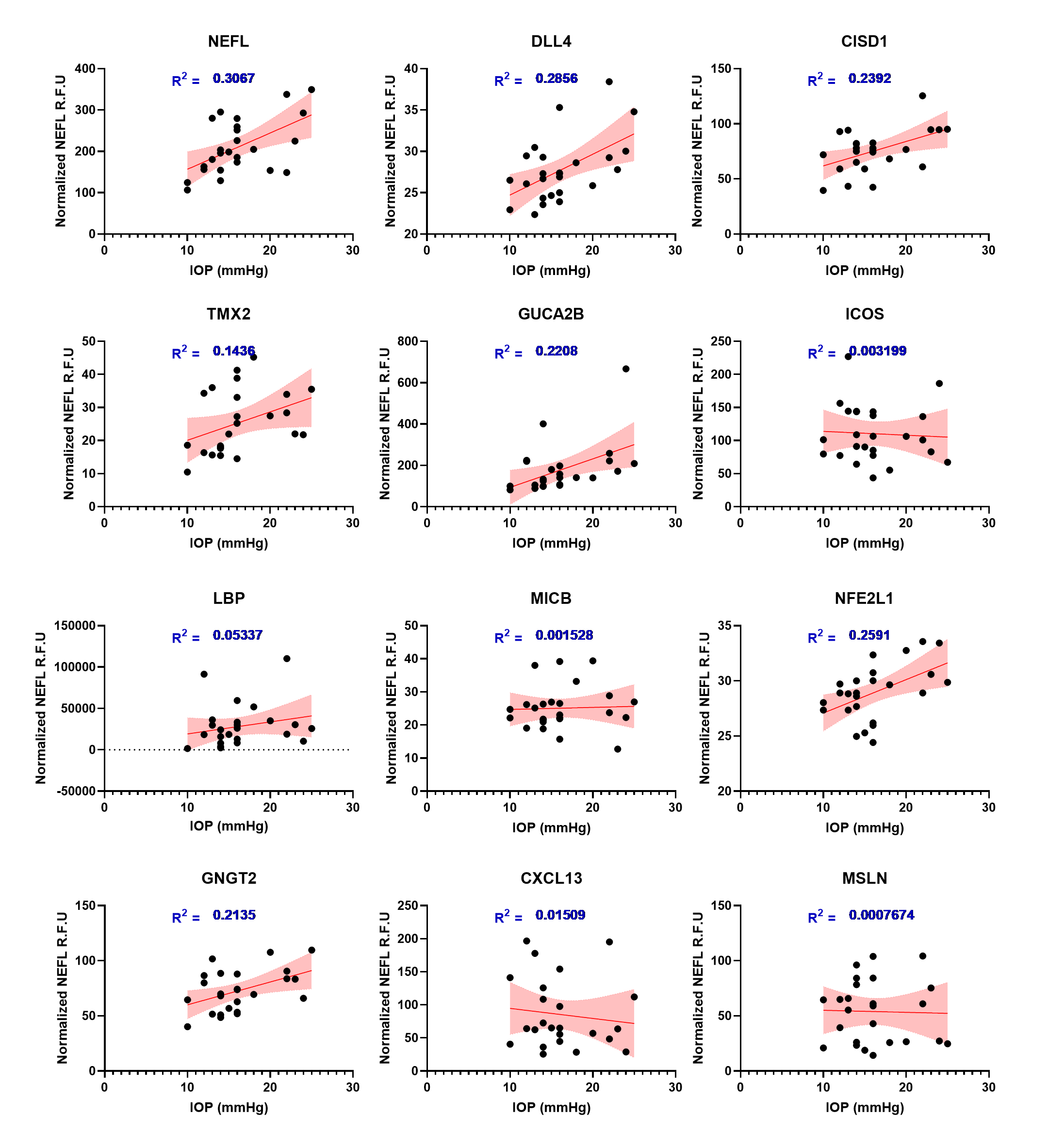

### Fig S3

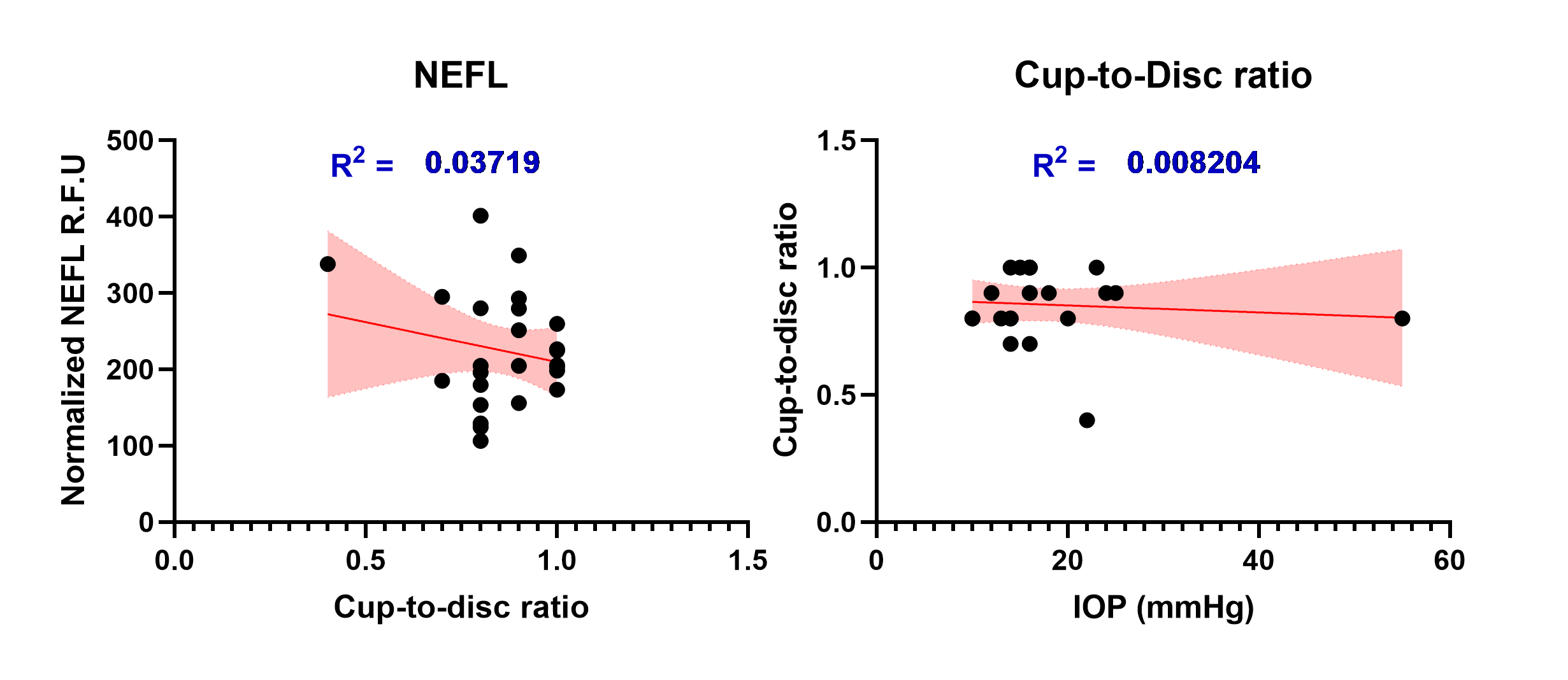
